# An aPKC rheostat induces apical contraction in response to epithelial stretching

**DOI:** 10.64898/2026.05.05.722904

**Authors:** Hélène Doerflinger, Amandine Palandri, Noah Jackaman, Ying Chen, Xixi Zhu, Daniel St Johnston

**Author notes:** Reproductive Medicine Center, The First Affiliated Hospital of Chongqing Medical University, Chongqing, China.

## Abstract

Apical-basal polarity in epithelial cells is controlled by a conserved set of polarity factors that define the apical, junctional and basolateral domains of the cell, but how these factors adapt to or control changes in domain sizes during cell shape changes remains unclear. Atypical protein kinase C (aPKC) is the main effector of apical identity, phosphorylating the lateral factors, Bazooka/Par-3, Lgl, Par-1 and Yurt to exclude them from the apical domain. Using analogue-sensitive aPKC in *Drosophila* follicle cells, we found that aPKC substrates differ over 100-fold in their sensitivity to inhibition, revealing a hierarchy of substrates, that is conserved in mammals, in which high-affinity substrates out-compete low-affinity substrates when aPKC activity is limiting. Mild aPKC inhibition prevents the phosphorylation of its lowest affinity substrate, Yurt. Yurt then accumulates apically by binding to Crumbs, where it activates apical constriction through Shroom, Cysts/Dp114RhoGEF, Rho kinase and Myosin. Yurt localises apically in cells that are stretched, either by morphogenesis or artificially, indicating that stretching reduces aPKC activity to trigger an antagonistic contraction. By contrast, *yurt*^-^ cells fail to resist stretching. Thus, the aPKC/Yurt pathway functions as a homeostatic stretch response, in which apical and lateral epithelial polarity factors collaborate to mechanically regulate apical domain size.

## Introduction

Many tissues and organs are composed of epithelial cells that adhere to each other to form sheets or tubes that function as barriers between compartments. The form and function of epithelia cells depends on their apical-basal polarisation, which directs the lateral localisation of adhesion molecules like E-cadherin that hold the epithelium together and targets different receptors, ion channels and transporters to their apical and basal membranes. This polarity is controlled by conserved proteins that define the apical, junctional and basolateral membrane domains[1]. Mutual antagonism between the apical and basolateral factors maintains this asymmetry by limiting the spread of each domain to position the apical junction. During development, epithelial cells can change shape to form squamous, cuboidal and columnar epithelia or undergo morphogenetic movements such as epithelial folding and epithelial to mesenchymal transitions. These processes require changes in the relative sizes of the apical, lateral and basal domains, but how polarity and morphogenesis are coupled is unclear[2].

aPKC is thought to be the main effector of apical identify, as all other apical factors are required to recruit it to the apical membrane and activate its kinase activity, where it plays a key role in defining epithelial polarity by excluding junctional and lateral polarity factors from the apical domain. aPKC phosphorylates Baz (Par-3) restrict it to the top of the lateral domain and position the apical junction [3–6]; and phosphorylates the lateral proteins, Lgl, Par-1 and Yurt, to prevent them from binding to the apical membrane [7–18].

Although these results indicate that the aPKC is essential for epithelial polarity, two reports have suggested that its kinase activity is not required in all contexts. Firstly, a single amino acid change in the activation loop of *Drosophila* aPKC that severely reduces its kinase activity in vitro has no detectable effect on apical-basal polarity in homozygous mutant follicle cells [19]. Secondly, *Drosophila* follicle cells treated with a specific inhibitors for either aPKC or Pak1 still exclude Lgl from the apical domain, whereas treatment with both inhibitors simultaneously leads to a mislocalisation of the apical factors and apical localisation of Lgl [20]. These results suggest that aPKC and Pak1 are redundant effectors of Cdc42 in specifying apical identity. This raises the question of why aPKC is required for polarity if its kinase activity is not essential.

We set out to confirm that aPKC’s kinase activity is dispensable, using an analogue-sensitive allele developed by Hannaford et al. [21]. Our results demonstrate that aPKC is not redundant with Pak1 and that its kinase activity is essential for polarity. More significantly, we observed that the phosphorylation of some aPKC substrates is more than two orders of magnitude more sensitive to inhibition of aPKC kinase activity than others, indicating that aPKC has a hierarchy of substrates, in which low affinity substrates are out-competed by high affinity substrates when the amount of active aPKC is limiting. Acutely blocking the phosphorylation of the lower affinity substrates reveals a novel role for aPKC in the control of apical constriction.

## Results

We first repeated the experiments that suggested that aPKC kinase activity is not essential using the follicle cells that surround developing *Drosophila* germline cysts as model. Previous work examined whether Lgl-GFP expressed from a UAS-transgene is excluded from the apical membrane of the follicle cells when aPKC is impaired or inhibited. It is possible that the Gal4 -mediated over-expression of Lgl-GFP alters the efficiency of its apical exclusion and we therefore assayed the apical exclusion of an endogenously-tagged version of Lgl, as well as a protein trap allele of a second aPKC substrate, Par-1 [22,23]. Both proteins are also expressed in the germline, however, and localise to the plasma membrane of the nurse cells and oocyte, which is hard to distinguish from the apical membrane of the follicle cells. We therefore also knocked down the expression of each protein in the germ line using nosGal4 to drive GFP-RNAi (Figs 1A and B).

**Figure 1.**
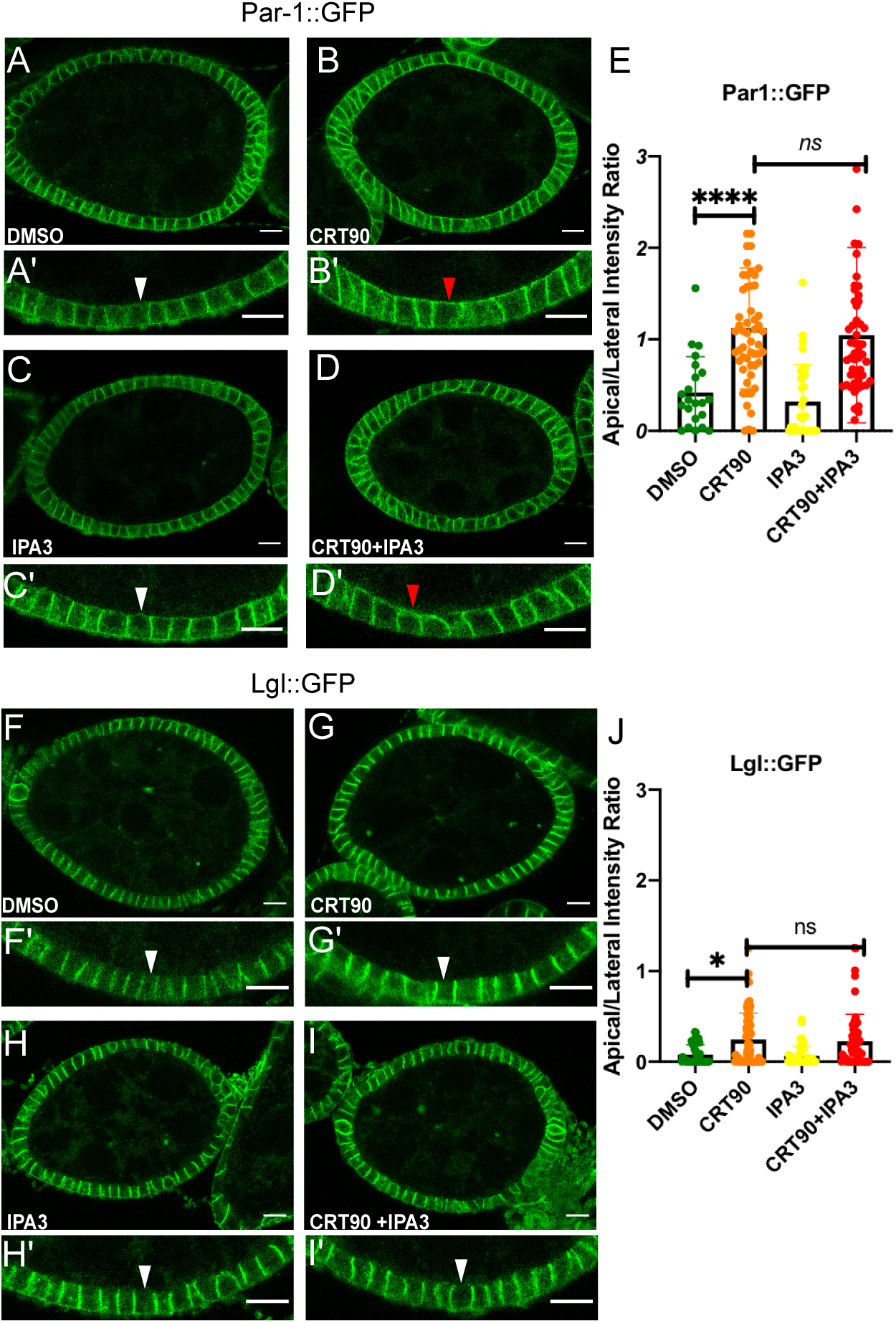
aPKC and Pak1 are not redundant Lgl kinases. **A-D**. Stage 6 egg chambers expressing endogenously-tagged GFP-Par-1. GFP-Par-1 was knocked down in the germ line by UAS-GFP RNAi driven by maternal a4 tubulin Gal4. **A.** Control egg chamber treated with DMSO for 1 hour. **A’.** A close up of the follicle cell layer in **A** showing the lateral localisation of GFP-Par-1. **B.** An egg chamber treated with 10µM of the aPKC inhibitor CRT90 in DMSO. GFP-Par-1 localises apically as well as laterally (red arrowhead in **B’**). **C** and **C’**. An egg chamber treated with 10µM of the Pak1 inhibitor IPA-3. GFP-Par-1 does not localises apically (white arrowhead in **C’**). **D** and **D’**. An egg chamber treated with 10µM IPA-3 and 10µM CRT90, showing the same apical localisation of Par-1 as with CRT90 alone (red arrowhead in **D’**). **E**. Quantification of the intensity of GFP-Par-1 signal at the apical membrane relative to the intensity at lateral membranes in control egg chambers treated with DMSO (n=22), egg chambers treated with CRT90 (n=55), with IPA-3 (n=33) and with IPA-3 and CRT90 (n=55). **F-I**. Stage 6 egg chambers expressing endogenously-tagged Lgl-GFP. Lgl-GFP was knocked down in the germ line by UAS-GFP RNAi driven by maternal ⍺4 tubulin Gal4. **F.** Control egg chamber treated with DMSO for 1 hour. **F’**. A close-up of the follicle cell layer in **F** showing the lateral localisation of Lgl-GFP. **G** and **G’**. An egg chamber treated with 10µM of the aPKC inhibitor CRT90 in DMSO. Lgl-GFP remains lateral (red arrowhead in **G’**). **H** and **H’**. An egg chamber treated with 10µM of the Pak1 inhibitor IPA-3. Lgl-GFP1 does not localises apically (white arrowhead in **H’**). **I** and **I’**, An egg chamber treated with 10µM IPA-3 and 10µM CRT90, showing the same lateral localisation of Lgl as with CRT90 alone (white arrowhead in **I’**). **J**. Quantification of the intensity of Lgl-GFP signal at the apical membrane relative to the intensity at lateral membranes in control egg chambers treated with DMSO (n=42), egg chambers treated with CRT90 (n=70), with IPA-3 (n=54) and with IPA-3 and CRT90 (n=59).

The evidence that aPKC and Pak1 are redundant comes from experiments in which egg chambers were treated with the specific aPKC inhibitors CRT0066854 or CRT0103390 (CRT90) and the Pak1 inhibitor, IPA-3 [20,24–26]. We therefore repeated these experiments using endogenously-tagged Par-1 and Lgl in egg chambers in which germ line expression of GFP was knocked down by RNAi. Treatment for 1 hour with the aPKC inhibitor CRT90 caused a significant increase in the level of GFP-Par-1 at the apical membrane compared to the DMSO control, whereas the Pak1 inhibitor had no effect (Figs 1A, 1B and 1E). Furthermore, treatment with both IPA-3 and CRT90 did not enhance the apical localisation of PAR-1 seen with CRT90 alone (Fig. 1C and 1D). By contrast, treatment with either or both inhibitors had no significant effect on the apical localisation of Lgl (Figs 1F, 1G, 1H, 1I, 1J). These results show that aPKC has a nonredundant function in excluding Par-1 from the apical domain but are consistent with the idea that its kinase activity may not be essential for the apical exclusion of Lgl. However, our data but do not support the proposal that aPKC is redundant with Pak1.

An alternative explanation for the lack of apical Lgl localisation treated with the aPKC inhibitor CRT90 is that it does not completely inhibit the kinase activity of aPKC and the residual activity is sufficient to exclude Lgl. We therefore took advantage of the analogue-sensitive aPKC allele, *aPKC*^as4^, in which the gate keeper mutation allows bulky non-hydrolysable ATP analogues to enter the ATP-binding pocket and inhibit kinase activity [21]. Treating wild-type egg chambers in culture with the cell-permeable, non-hydrolysable ATP analogue 1NA-PP1 has no effect on the localisation of Par-1 or Lgl, indicating that the analogue does not have off target effects (Fig.2A). By contrast, both Par-1 and Lgl relocalise to the apical membrane within 20 minutes of adding 1NA-PP1 to *aPKC*^as4^ homozygous egg chambers (Figs 2A and 2C). However, the concentration of the inhibitor required to induce apical localisation of each protein is different: Par-1 is fully apical after treatment with 1µm 1NA-PP1, whereas Lgl is only partially apical after treatment with 5µm 1NA-PP1 and fully apical after treatment with 10µm (Figs 2A, 2B, 2C and 2D).

**Fig. 2:**
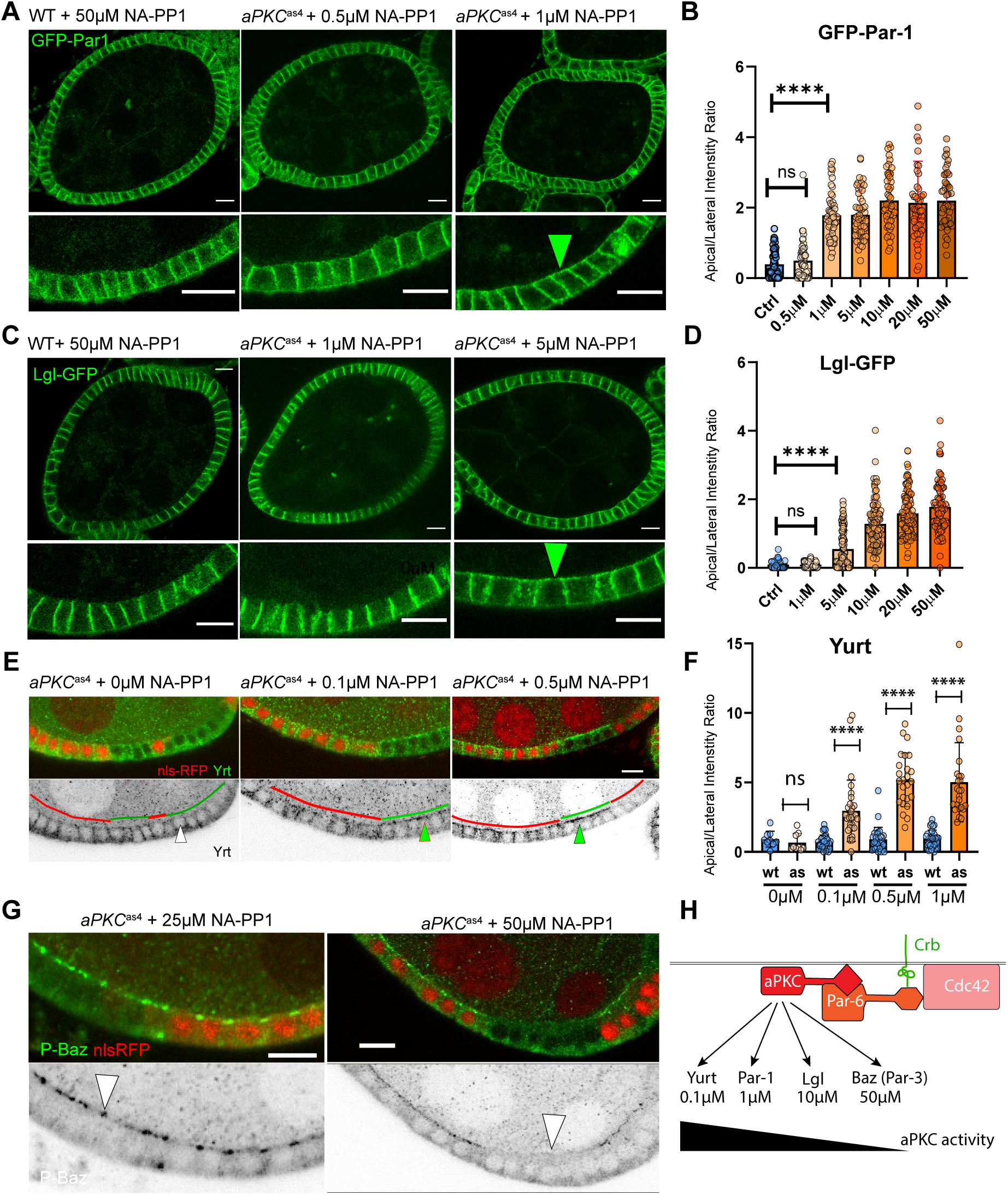
aPKC substrates differ 500-fold in the concentration of aPKC inhibitor that prevents their phosphorylation. **A,** Control and *aPKC*^as4^ egg chambers expressing endogenously-tagged GFP-Par-1 (green) treated for 20 min with various concentrations 1NA-PP1 (NA-PP1). The green arrowhead marks apical GFP-Par-1 after treatment with 1 µM 1NA-PP1. **B**, Quantification of the ratio of apical to lateral GFP-Par-1 intensity (n = 50, except 50µM, n = 46. **C**, Control and aPKC^as4^ egg chambers expressing endogenously-tagged Lgl-GFP treated with 1NA-PP1. GFP-Par-1 and Lgl-GFP were knocked down in the germ line by maternal a4 tubulin-Gal4 driven UAS-GFP RNAi in **A** and **C**. **D**, Quantification of the ratio of apical to lateral Lgl-GFP intensity (n = 90, 100, 100, 95, 100, 82). **E.** Egg chambers containing aPKC^as4^ homozygous clones marked by the loss of nls-RFP (red) stained for Yurt (green; black in lower panels). **F**, Quantification of the ratio of apical to lateral Yurt localisation in *aPKC*^as4^/+ control cells and *aPKC*^as4^ cells (0µM, n = 9, 9; 0.1µM, n = 31, 29; 0.5µM, n = 30, 25; 1µM, n = 30, 25). **G**, Egg chambers containing *aPKC*^as4^ homozygous clones marked by the loss of nls-RFP (red, marked by white arrowheads) treated with 1NA-PP1 and stained for phospho-BazS980 (25µM, n = 26; 50µM, n = 22). **H**, Model showing the hierarchy of aPKC substrates. Scale bars, 10 µm.

The tenfold difference in the concentration of 1NA-PP1 require to inhibit the apical exclusion of Par-1 and Lgl prompted us to examine how much inhibitor is required to block the phosphorylation of other aPKC substrates. To examine Yurt, which is also excluded from the apical domain by aPKC phosphorylation, we generated homozygous aPKC^as^ clones marked by the loss of RFP, so that the surrounding heterozygous cells provide an internal control for the antibody staining. Yurt already appears at the apical membrane after 20 minutes of exposure to 0.1µm 1NA-PP1 and full apical localisation occurs at 1µm (Figs 2E and 2F). By contrast, Bazooka is still phosphorylated at 25 µm 1NA-PP1 as detected by an antibody that recognises phosphorylated serine 980 of Bazooka (Fig. 2G) [3]. However, this phosphorylation is lost after treatment with 50µm 1NA-PP1 (Fig. 2G). There is therefore a 500-fold difference in the concentration of 1NA-PP1 required to inhibit aPKC phosphorylation of its most sensitive substrate (Yurt) and the least sensitive, Bazooka/Par-3.

These results demonstrate that aPKC is the sole kinase that acts to exclude lateral polarity factors from the apical membrane, suggesting that CRT90 only partially inhibits aPKC activity, so that only the most sensitive substrates are affected. In support of this view, 10µM CRT90 induced the rapid apical localisation of Yurt, consistent with Yurt’s position as the substrate that is most sensitive to aPKC inhibition (Supplementary Video 1). CRT90 therefore provides a useful tool for mildly inhibiting aPKC activity, without affecting most substrates or disrupting apical-basal polarity.

The large differences in the sensitivity of aPKC substrates to the aPKC inhibitor that we observed fit very well with in vitro studies with the catalytic domain of human aPKC, which demonstrated that aPKC substrates have very different affinities for the kinase, with Par-3 and Lgl binding more strongly (K_M_ = 0.24µM and 0.35µM) than Par-1 (K_M_ = 16.6µM) [27]. In conjunction with our data, these observations suggest that the high affinity aPKC substrates out compete lower affinity substrates when aPKC activity is limiting, giving rise to the hierarchy of substrates whose phosphorylation is blocked at different thresholds of aPKC inhibition (Fig. 2H). In support of this view, over-expression of the high affinity substrate, Lgl, led to the apical localisation of the low affinity substrate, Yurt, indicating that Yurt was not being phosphorylated by aPKC, even though the latter’s activity was unperturbed (Figs 3A and 3B).

**Fig. 3.**
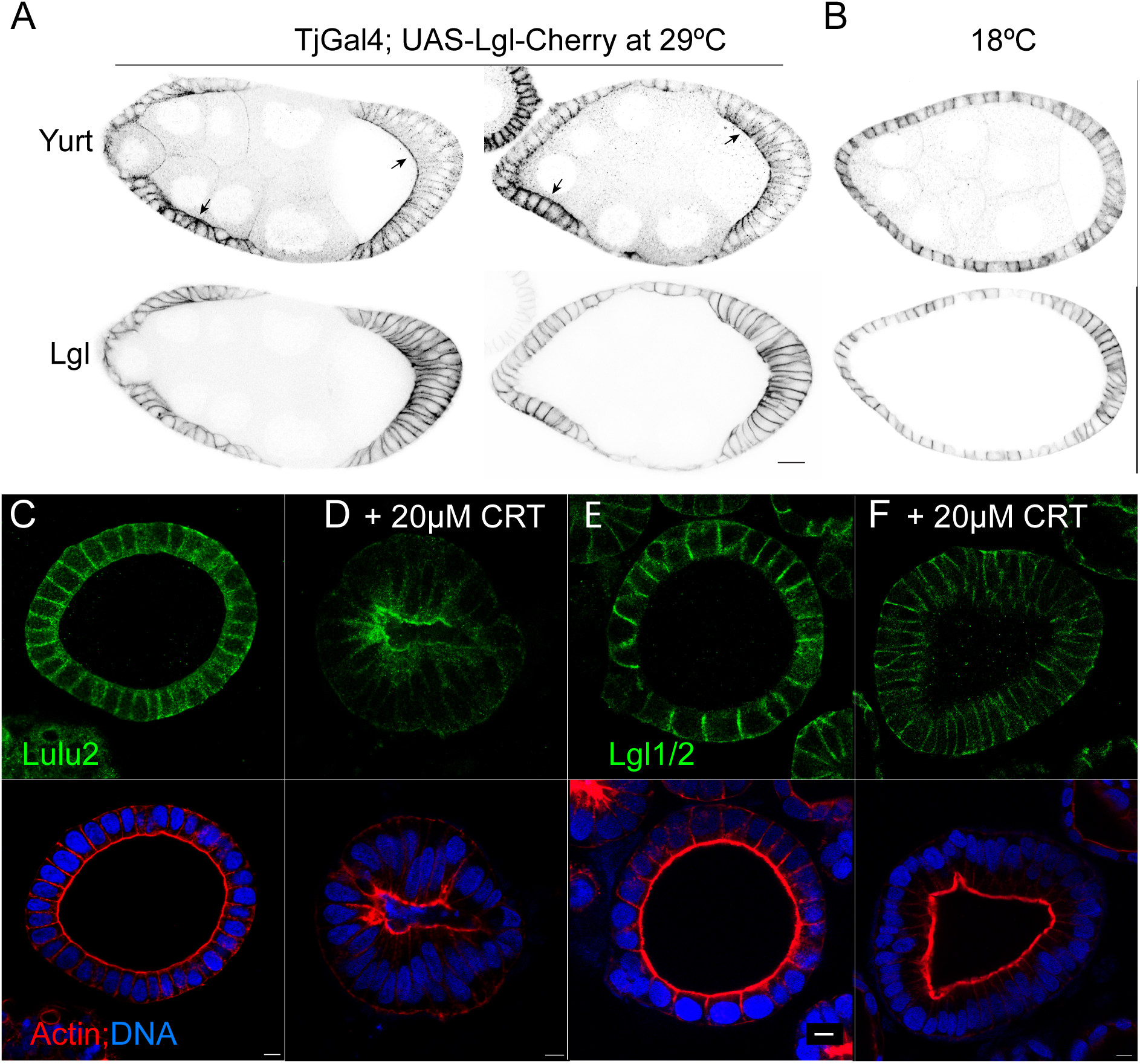
The hierarch of aPKC substrates is conserved in humans. **A-B**. Egg chambers expressing Lgl-Cherry under the control of Tj-Gal4 at 29_°_C (**A,** high expression) or 18_°_C (**B**, very low expression) stained for Yurt. High levels of Lgl outcompete Yurt for phosphorylation by aPKC, leading to the apical localisation of unphosphorylated Yurt, whereas Lgl remains lateral even when over-expressed. **C-D**, Caco-2 cysts in 3D culture stained for the Yurt orthologue, Lulu2 (green), Actin, (red) and DNA (blue). Lulu2 is lateral in the untreated cyst (**C**) but localises apically in the cyst treated with 20 µM CRT90 (**D). D-E**, Caco-2 cysts in 3D culture stained for Lgl1/2 (green), Actin, (red) and DNA (blue). Lgl1/2 localise laterally in both the untreated cyst (**D**) and the cyst treated with 20 µM CRT90 (**E**). Scale bars, 10µm.

Since human aPKC substrates also have different affinities for the kinase[27], we took advantage of the weak aPKC inhibitor, CRT90, to test whether there is a similar hierarchy of aPKC substrates in human epithelia. Untreated Caco-2 cells grown in 3D culture form epithelial cysts with an actin-rich apical domain facing an internal lumen (Figs 3C and 3D). The lateral polarity factors, Lulu2 (also called Epb41L4B/EHM2), an orthologue of Yurt, and Lgl1 and 2 localise to the lateral membrane and are excluded from the apical domain by aPKC[8,17]. Treatment of these cysts with 20µM CRT90 induced the apical localisation of Lulu2, but had no effect on the lateral localisation of Lgl1/2 (Figs 3C and 3D). Thus, Lulu 2 is more sensitive to aPKC inhibition than Lgl, just as in *Drosophila*, indicating that the hierarchy of aPKC substrates is conserved.

### aPKC inhibition induces apical constriction

These results raise the question of whether the hierarchy of aPKC substrates has biological significance. Treating egg chambers containing *aPKC*^as4^ homozygous clones with a low concentration of 1NA-PP1 induced apical constriction of the *aPKC*^as4^ homozygous cells, reducing the size of their apical domains and increasing their height (Figs 4A and 4B). Caco-2 cells showed the same response to partial aPKC inhibition, with the CRT90-treated cells becoming more columnar with smaller apical domains, indicating that partial aPKC inhibition also triggers apical constriction in mammalian cells (Figs 3C and 3D). Similarly, treating entirely *aPKC*^as4^ homozygous egg chambers with 1 µM 1NA-PP1 or wild-type egg chambers with 10 µM CRT90 caused the egg chambers to become rounder as the follicle cells apically constrict around the incompressible germline cyst (Supplementary Video 2). Inhibiting aPKC with 10µM CRT90 in egg chambers expressing the Myosin regulatory light chain-GFP (MRLC-GFP) revealed that this apical constriction correlates with the apical enrichment of Myosin (Figs 4C and C’; Supplementary Video 3). Thus, reducing aPKC activity to a level that only inhibits the phosphorylation of its lowest affinity substrates increases apical tension, in good agreement with the observation that optogenetic inhibition of aPKC induces apical constriction and can rupture the follicular epithelium when cells enter mitosis[28].

**Fig 4.**
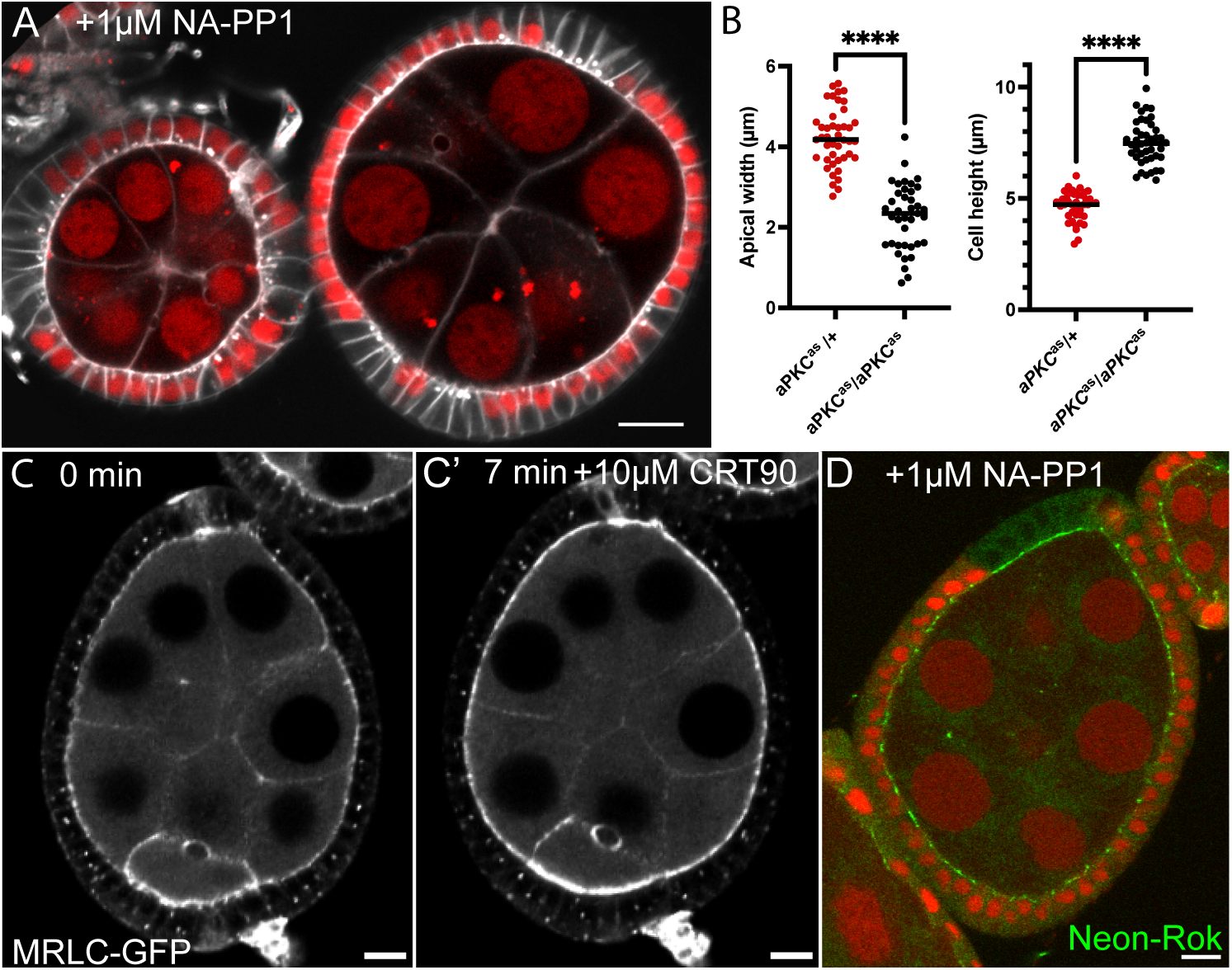
Reduced aPKC activity induces apical constriction and the apical localisation of Myosin and Rho kinase. **A**. Egg chambers containing aPKC^as4^ homozygous clones marked by the loss of nls-RFP (red) and treated with 2 µM 1NA-PP1. The aPKC^as4^ homozygous cells apically constrict, reducing the size of their apical domains and increasing cell height. **A’**. Quantification of the shape change of *aPKC*^as4^ cells after 1NA-PP1 treatment compared to their heterozygous neighbours. **B**. An *aPKC*^as4^ egg chamber expressing the Myosin regulatory light chain fused to GFP (MRLC-GFP) before and 7 minutes after treatment with 1 µM 1NA-PP1. The partial inhibition of aPKC induces apical accumulation of Myosin and apical constriction, making the egg chambers rounder. **C**. An egg chamber expressing Neon-Rok (green) containing aPKC^as4^ homozygous clones marked by the loss of nls-RFP (red) and treated with 1 µM 1NA-PP1. Neon Rok localises to the apical junctions in the heterozygous *aPKC*^as4^/+ cells, but accumulates apically in the homozygous cells. Scale bars, 10µm.

During many morphogenetic processes involving apical constriction, such as *Drosophila* ventral furrow formation or salivary placode invagination, Myosin is activated by Rho-kinase [29–31]. We therefore examined the localisation of endogenously-tagged Neon-Rok in *aPKC*^as4^ homozygous clones treated with 1µM 1NA-PP1 (Fig. 4D). Rok became highly enriched at the apical surface of the *aPKC*^as4^ cells, whereas it localised to the adherens junctions at the apical/lateral borders of the control heterozygous cells. When these clones were imaged in living egg chambers ex vivo, partial aPKC inhibition induced a marked apica enrichment of Rok in the contracting *aPKC*^as4^ homozygous cells (Supplementary Video 4).

### Yurt activates the Cysts/Rho kinase pathway

The apical constriction of the follicle cells when aPKC is partially inhibited is presumably caused by the loss of phosphorylation of a low affinity aPKC substrate. A strong candidate for this substrate is Yurt, since it lies at the bottom of the hierarchy of aPKC substrates and because the over-expression of both *Drosophila* Yurt and its mammalian orthologues, Lulu 1 and 2 cause apical constriction in epithelial cells[32,33]. We therefore induced apical constriction using CRT90 in Neon-Rok-expressing egg chambers that also contained *yurt* null mutant clones (Supplementary video 5). The *yurt* mutant cells had the same shape and as the wild-type cells before the addition of CRT90 (Fig. 5A). After CRT90 addition, the wild-type cells accumulated high levels of apical Rok and apically constricted as expected (Fig. 5B). By contrast, the *yurt* mutant cells did not accumulate apical Rok or constrict, and instead they became stretched by the contraction of their neighbours. Thus, mild aPKC inhibition induces apical constriction by preventing the phosphorylation of Yurt, which then localises apically, leading to the activation of Rok and Myosin. The inability of *yurt* mutant cells to resist stretching may account for the elongated phenotype of a significant proportion of *yurt* mutant clones in untreated egg chambers, suggesting that these cells have been stretched by mechanical forces during normal development.

**Fig 5.**
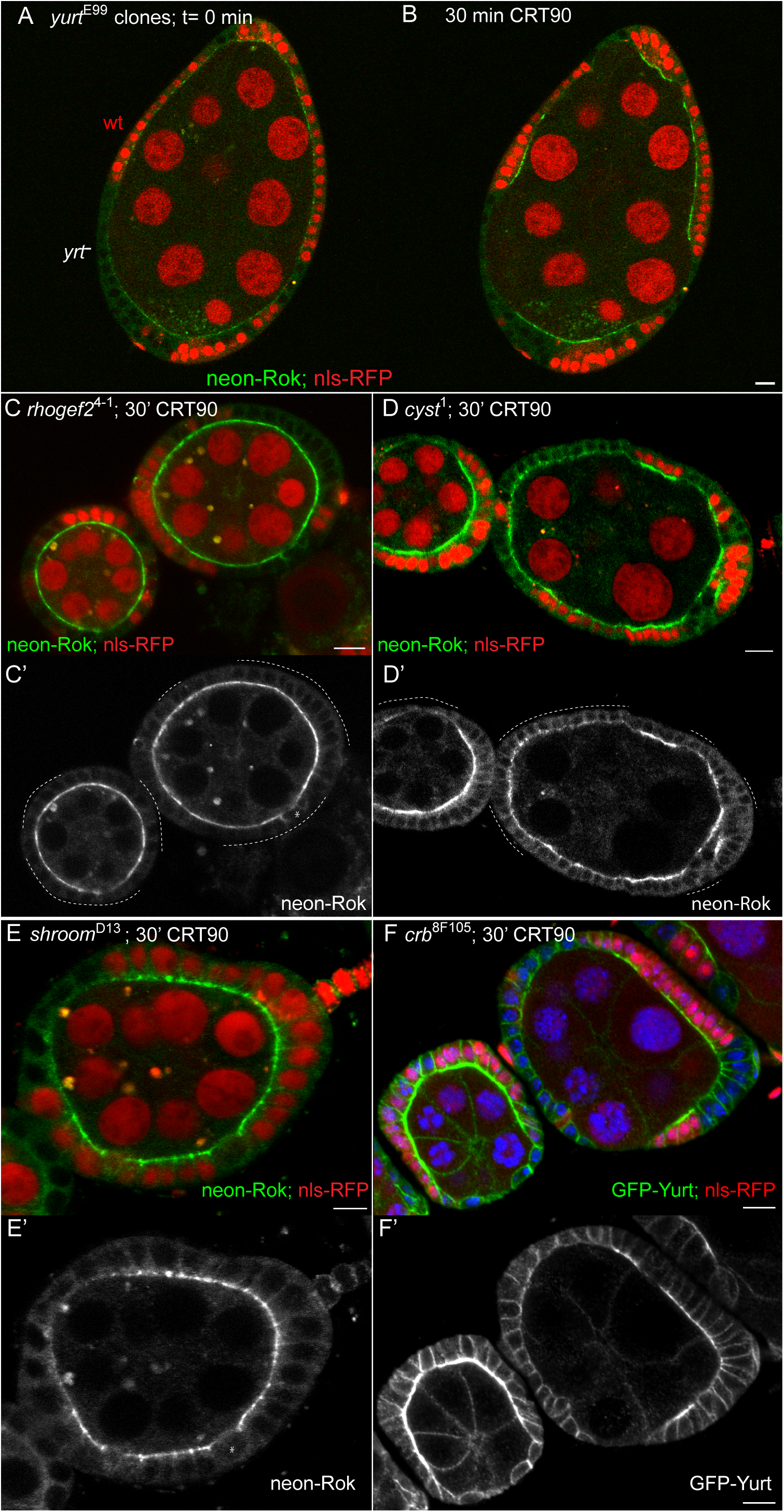
Mild aPKC inhibition activates junctional contractility through Yurt, Crumbs, Shroom, Cysts and Rok. **A**, An egg chamber expressing Neon-Rok (green) containing *yurt*^E55^ clones marked by the loss of nls-RFP (red) before the addition of CRT90. The *yurt* mutant cells are similar in shape to their heterozygous neighbours. **B**. The same egg chamber 30 minutes after addition of 10 µM CRT90. The heterozygous cells become more columnar and localise Neon-Rok apically after mild aPKC inhibition, whereas the *yurt* mutant cells do not localise Neon-Rok and become stretched by the contraction of their neighbours. **C** and **C’.** An egg chamber expressing Neon-Rok (green; white in **C’**) containing *rhogef2*^1^ clones marked by the loss of nls-RFP (red) after treatment with 10 µM CRT90. The localise Neon-Rok localises apically after partial aPKC inhibition in both *rhogef2* mutant cells and their heterozygous neighbours. The white asterisk marks a cell in mitosis when apical markers delocalise. **D** and **D’**. An egg chamber expressing Neon-Rok (green; white in **D’**) containing *cysts*^1^ clones (a.k.a. *Dp144rhogef*) marked by the loss of nls-RFP (red) after treatment with 10 µM CRT90. The *cysts* mutant cells show little or no apical localisation of Neon-Rok on aPKC inhibition compared to their heterozygous neighbours and remain cuboidal. **E** and **E’**, An egg chamber expressing Neon-Rok (green; white in **E’**) containing *shroom*^1^ clones marked by the loss of nls-RFP (red) after treatment with 10 µM CRT90. The *shroom* mutant cells fail to localise Neon-Rok apically after aPKC inhibition and are shorter than the heterozygous cells. The white asterisk marks a cell in mitosis. **F** and **F’**. An egg chamber expressing GFP-Yurt (green; white in **F’**) containing *crb*^F105^ clones marked by the loss of nls-RFP (red) after treatment with 10 µM CRT90. The *crb*^-^ cells do not localise GFP-Yurt apically after mild aPKC inhibition, and many cells become shorter and stretched. Scale bars, 10 µm, except **E**, which is 5 µm.

Rok is normally activated apically by Rho1, which is in turn activated by different Rho guanine nucleotide exchange factors (RhoGEFs) at different apical positions. RhoGEF2 activates Rho at the medial-apical cortex, during ventral furrow formation for example, whereas Cysts/p114RhoGEF activates Rho at the apical adherens junctions to control junctional tension[34–36]. To test whether either RhoGEF is required for the apical recruitment of Rok after partial aPKC inhibition, we generated *rhogef2* and *cysts* mutant clones in Neon-Rok expressing egg chambers treated with CRT90 (Figs 5C and D). Loss of RhoGEF2 had no effect on the apical accumulation of Rok on aPKC inhibition, whereas the *cysts* mutant cells had much lower levels of apical Rok than their wild-type neighbours, and were shorter and more stretched, indicating that they were not undergoing apical constriction. This is consistent with the observation that a human Yurt orthologue, Lulu2 binds to p114RhoGEF directly, suggesting that apical Yurt recruits Cysts to activate Rho and Rok to induce myosin contractility[37]. Like Yurt and Lulu1/2, the Shroom family of proteins induce apical constriction, in this case by recruiting Rok directly, and this activity depends on Lulu in *Xenopus*[38–42]. We therefore tested whether the single *Drosophila* Shroom is involved in Yurt-dependent apical constriction. Indeed, *shroom* mutant clones failed to accumulate Neon-Rok apically after treatment with CRT90 and became shorter than their wild-type neighbours (Figs 5E and 5E’). Finally, we tested the role of Crumbs, since Yurt binds directly to the FERM-binding domain of Crumbs[43,44]. *crb* mutant follicle cells in which aPKC had been inhibited with CRT90 also failed to constrict and became stretched due to the contraction of their neighbours (Figs 5F and 5F’). Yurt still localised weakly to the apical domain in the *crb* mutant cells, because it binds the plasma membrane when not phosphorylated by aPKC, but does not become highly enriched there, since it cannot bind Crumbs and tetramerise[18,45]. Thus, by allowing the apical Yurt accumulation in association with Crumbs, reduced aPKC activity triggers the junctional contractility pathway through Cysts, Shroom, Rok and Myosin.

### The aPKC/Yurt pathway acts as a stretch response

The observation that *yurt* mutant cells become abnormally stretched when pulled on by their neighbours suggests that the aPKC/Yurt contractility pathway plays a role in epithelial cells’ response to stretching. To test this hypothesis, we examined the follicle cells during the 40 minute transition from stage 10b to 11, when the nurse cells rapidly dump their contents into the oocyte, leading the oocyte to nearly double in length, which stretches the follicle cells that surround it[46]. Yurt is exclusively lateral in stage 10b follicle cells, but becomes enriched apically at stage 11 in the stretched follicle cells (Figs 6A and 6B). This apical enrichment becomes even more pronounced at stage 12 when the oocyte reaches its final size and the follicle cells are even more elongated (Fig. 6C). While Yurt becomes exclusively apical in the follicle cells along the sides of the oocyte, the follicle cells around the oocyte posterior show a weaker apical localisation of Yurt, with residual Yurt along the lateral cortex and a pronounced enrichment at the apical adherens junctions (Figs 6C’ and 6C”). These cells are less stretched than the other follicle cells, because the oocyte mainly expands along its anterior-posterior axis due to the molecular corset of circumferential collagen fibres established by egg chamber rotation[47]. This suggests that the response to stretching has two phases, in which Yurt localises to the adherens junctions after mild stretching and then extends across the entire apical domain as the cells become more elongated. To test whether these phases of Yurt localisation correlate with the degree of aPKC inhibition, we treated GFP-Yurt-expressing egg chambers with a 5-fold lower concentration of CRT90 (2µM) to reduce aPKC activity more gradually (Supplementary video 6). Yurt first became enriched at the adherens junctions of the follicle cells in treated egg chambers, before spreading apically after 20 minutes. Thus, the level of the reduction in aPKC activity determines whether Yurt is recruited to adherens junctions only or to the entire apical domain.

**Fig 6.**
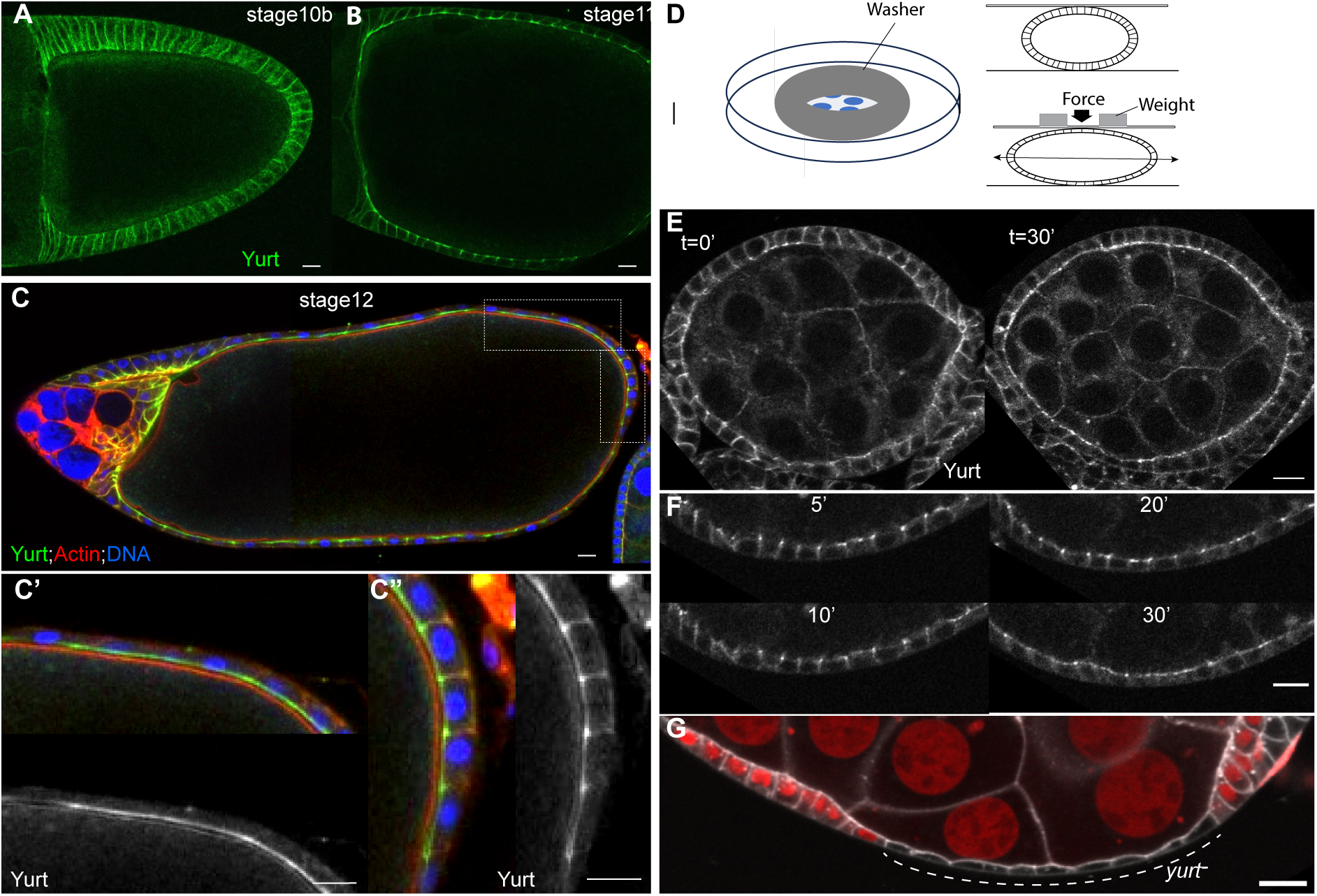
Epithelial cell stretching induces apical Yurt localisation. **A**. Endogenously-tagged GFP-Yurt localises laterally in the follicle cells of a stage 10b egg chamber, before nurse cell dumping has begun. **B.** GFP-Yurt localises apically in a stage 11 egg chamber part way through dumping, when the follicle cells are stretched. **C-C”.** A stage 12 egg chamber near the end of nurse cell dumping, expressing GFP-Yurt (green) and stained for F-actin (red) and DNA (blue). Yurt localises entirely apically in the follicle cells along the length of the oocyte (**C’**), but is still partly lateral and enriched at the apical junctions of cells at the posterior pole, which are less elongated (C”). **D.** A drawing showing the design of the egg chamber compression experiments. **E.** A stage 5 egg chamber expressing endogenously-tagged GFP-Yurt before (left), and 30 minutes after compression (right). Yurt localises to the apical side of the follicle cells after compression. **F.** Time points from the compression of another egg chamber, showing that Yurt accumulates first at apical junctions (10’), before spreading across the rest of the apical domain (20’and 30’). **G.** A compressed egg chamber containing a *yurt*^E99^ clone marked by the loss of nls-RFP (red) and labelled with CellMask. The *yurt* mutant cells do not resist stretching and become more elongated than the heterozygous cells. Scale bars 20 µm (**A**-**C**) and 10 µm (**E**-**G**).

Since the elongation of the follicle cells at stage 11-12 is a normal developmental process, it is possible that the apical accumulation of Yurt is genetically programmed rather than a response to stretching. To rule out this possibility, we mechanically stretched early, Yurt-GFP expressing egg chambers that do not normally elongate by culturing them ex vivo and applying a smally weight to the coverslip above them (Fig. 6D). This led to the complete relocalisation of Yurt from the lateral to the apical side of the follicle cells in 30 minutes (Supplementary videos 7-8; Fig. 6E). A time course of a compressed egg chamber showed that Yurt first accumulated at the Adherens junctions before spreading across the apical surface, consistent with two-phase response to a reduction in aPKC activity (Fig. 6F; Supplementary video 9). Epithelial cell stretching therefore induces the junctional/apical localisation of Yurt, thereby activating Myosin to produce a contraction force that counteracts the stretch, providing a homeostatic mechanism to maintain apical domain size. Consistent with this, when egg chambers containing *yurt* clones were compressed, the *yurt* mutant cells became highly elongated compared to their wild-type neighbours, indicating that Yurt is essential for epithelial cell resistance to stretching (Fig. 6G). Yurt is often slightly enriched at the apical junctions of wild-type follicle cells at earlier stages of oogenesis, and this enrichment is lost in *crb* mutant cells (Figure S1). This suggests that follicle cells are normally subject to some degree of stretching and that the level of aPKC activity is set close to the threshold for Yurt apical localisation. This ability of epithelial cells to resist stretching is likely to be particularly important during development when morphogenetic movements exert pulling forces on neighbouring epithelia.

## Discussion

The response of epithelial cells to stretching in tissue culture models has led to the discovery that both Piezo and the YAP/TAZ pathway respond to stretching by activating cell division to restore normal cell density[48–51]. These are slow processes that can take hours to days, however, and recent work has identified a much more rapid response in vivo, in which stretched epithelial cells localise contractile actomyosin apically within minutes, but how this occurs is not understood [52,53]. Our results showing that epithelial cell stretching reduces aPKC activity to activate junctional contractility are likely to provide the mechanism that underlies this fast response.

One question raised by our results is how apical domain expansion leads to a reduction in aPKC activity to allow Yurt to localise apically. One possibility is that upstream factors, such as stretch-activated ion channels, somehow induce a decrease in aPKC activity when the apical domain expands[54,55]. However, a simpler model is that stretching the apical domain directly reduces aPKC activity (Fig. 7). aPKC is only fully active when it is localised at the apical plasma membrane, since its activation depends on its binding to Par-6, the binding of Par-6 to Cdc42-GTP, which is exclusively apical and the binding of the Par-6 PDZ domain to the apical transmembrane protein Crumbs[56–59]. This means that active aPKC will be diluted when the apical domain expands, whereas concentrations of its cytoplasmic substrates will not change. Apical domain expansion could therefore cause the density of active aPKC at the membrane to fall below the threshold required to phosphorylate its lowest affinity substrate, Yurt, because of competition with higher affinity substrates, like Lgl. This proposed mechanism differs from other mechano-sensitive responses in that it couples myosin activation to the change in apical domain size rather than the stretching force itself [60,61].

**Figure 7.**
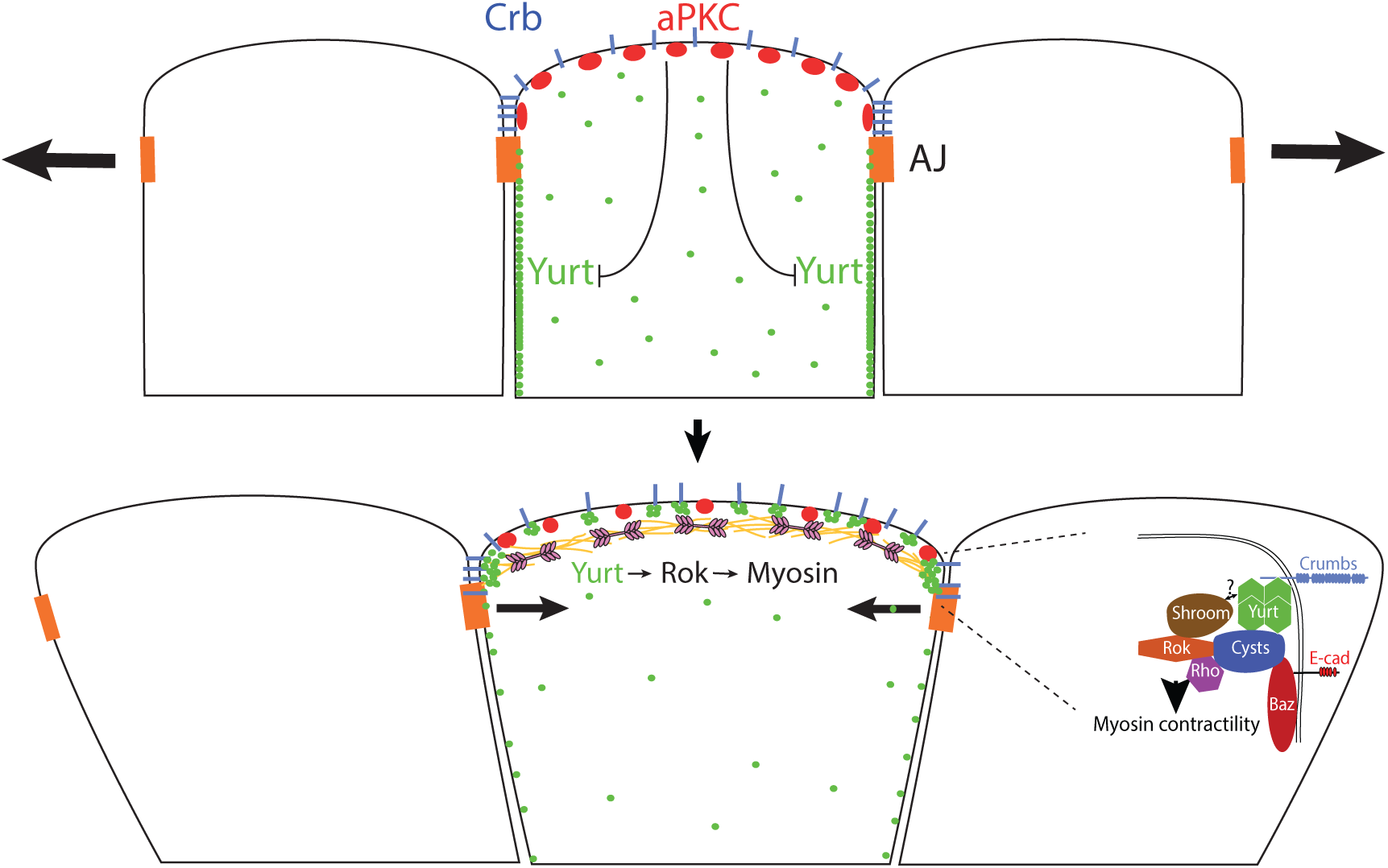
A model showing how cell stretching could activate a compensatory apical contraction. Apical domain expansion dilutes active aPKC at the apical membrane, allowing unphosphorylated Yurt to bind to Crumbs and tetramerise, where it induces Myosin activation through Shroom, Cysts, Rho, Rho kinase and Myosin.

Whatever the mechanism, the stretch-induced reduction in aPKC activity turns on junctional contractility by allowing Yurt to localise apically by binding to Crumbs. Yurt then acts through Cysts/p114RhoGEF, Shroom and Rho kinase to activate Myosin contractility. All of the proteins downstream of Crumbs and Yurt are components of the junctional contractility pathway that drives multiple morphogenetic movements, such as germband extension in *Drosophila*, but the mechanisms that activate this pathway are not well understood. It will therefore be important to investigate whether the modulation of aPKC activity and apical Yurt recruitment by Crumbs provide a general mechanism to control this pathway in a variety of contexts.

The relative sizes of the apical and lateral domains of epithelial cells are thought to be determined by activity of the polarity proteins in each domain and the mutual antagonism between them. In this view, Yurt has been considered a lateral polarity factor that directly inhibits the function of the apical polarity factor, Crumbs, because Yurt can bind to Crumbs and because *yurt* mutant epithelial cells show expanded apical domains, a phenotype that mimics the effects of Crumbs over-expression[43,62]. Our results and those of Biehler et al.[32] suggest an alternative explanation for these observations, in which loss of Yurt increases apical domain size through the loss of junctional tension, rather than by antagonising Crumbs. In fact, the binding of Yurt to Crumbs is necessary for the apical recruitment of Rok and Myosin and the activation of contractility. Thus, rather than inhibiting Crumbs, Yurt functions with Crumbs to control cell mechanics. This role of Crumbs can explain the observation that it is only required in embryonic tissues that are undergoing morphogenesis[63]. Like Yurt, Lgl over-expression decreases the size of the apical domain and increases that of the lateral domain[64,65]. This is also probably due to apical constriction, as over-expressed Lgl out-competes Yurt for phosphorylation by aPKC, allowing Yurt to localise apically. Thus, much of the observed antagonism between apical and lateral polarity proteins can be attributed to mechanical effects rather than inhibition, demonstrating that the polarity system directly regulates cell shape.

The function of aPKC as a stretch sensor depends on its behaviour as a rheostat that phosphorylates fewer substrates as its activity decreases. This unusual property is shared with two other key regulatory kinases, TORC1 and Cdk1. TORC1 phosphorylates substrates that activate different biosynthetic pathways at different thresholds, and modulation of Cdk1 activity is sufficient to drive the entire *S. pombe* cell cycle: low levels of active Cdk phosphorylate the substrates that initiate S phase, and higher levels of activity are required to phosphorylate the substrates that trigger mitosis[66,67]. The 100-500 fold differences in the sensitivity of aPKC substrates to kinase inhibition seem to depend on the affinity of each substrate to the active site of aPKC. The ability of the high affinity substrates to outcompete lower affinity substrates for binding to the active site of aPKC is further enhanced by the low turn-over rate of the high affinity substrates, Par-3 and Lgl, which means that they remain bound to the active site block the active site for longer, thereby preventing other substrates from binding [27]. Furthermore, both Baz/Par3 and Lgl also act to inhibit aPKC kinase activity up binding [68,69]. Thus, they should be very effective at outcompeting lower affinity substrates when aPKC activity is limiting. Although the hierarchy of aPKC substrates is most likely determined by the binding affinity of each substrate for the aPKC active site, it is also possible that the activity of the phosphatases that dephosphorylate the aPKC sites in each substrate also contributes.

Yurt is the first well-characterised substrate that is not phosphorylated as aPKC activity decreases, and the threshold for this switch defines the sensitivity of the stretch response pathway. Above this threshold, the level of aPKC activity determines the amount of unphosphorylated Yurt and whether Yurt localises just to Adherens junctions, where Crumbs is concentrated, or across the whole apical domain. Since some Yurt accumulates to at the apical junction of many wild-type cells, the stretch response pathway appears to be often activated at low levels, suggesting that cells in the epithelium use this mechanism to constantly respond to changes in apical domain size. In this way, aPKC can act both as a switch and a rheostat to regulate the level of Myosin contractility as the apical domain expands, providing an elegant mechanism to produce a rapid and proportional response to cell stretching.

## Materials and Methods

## Materials

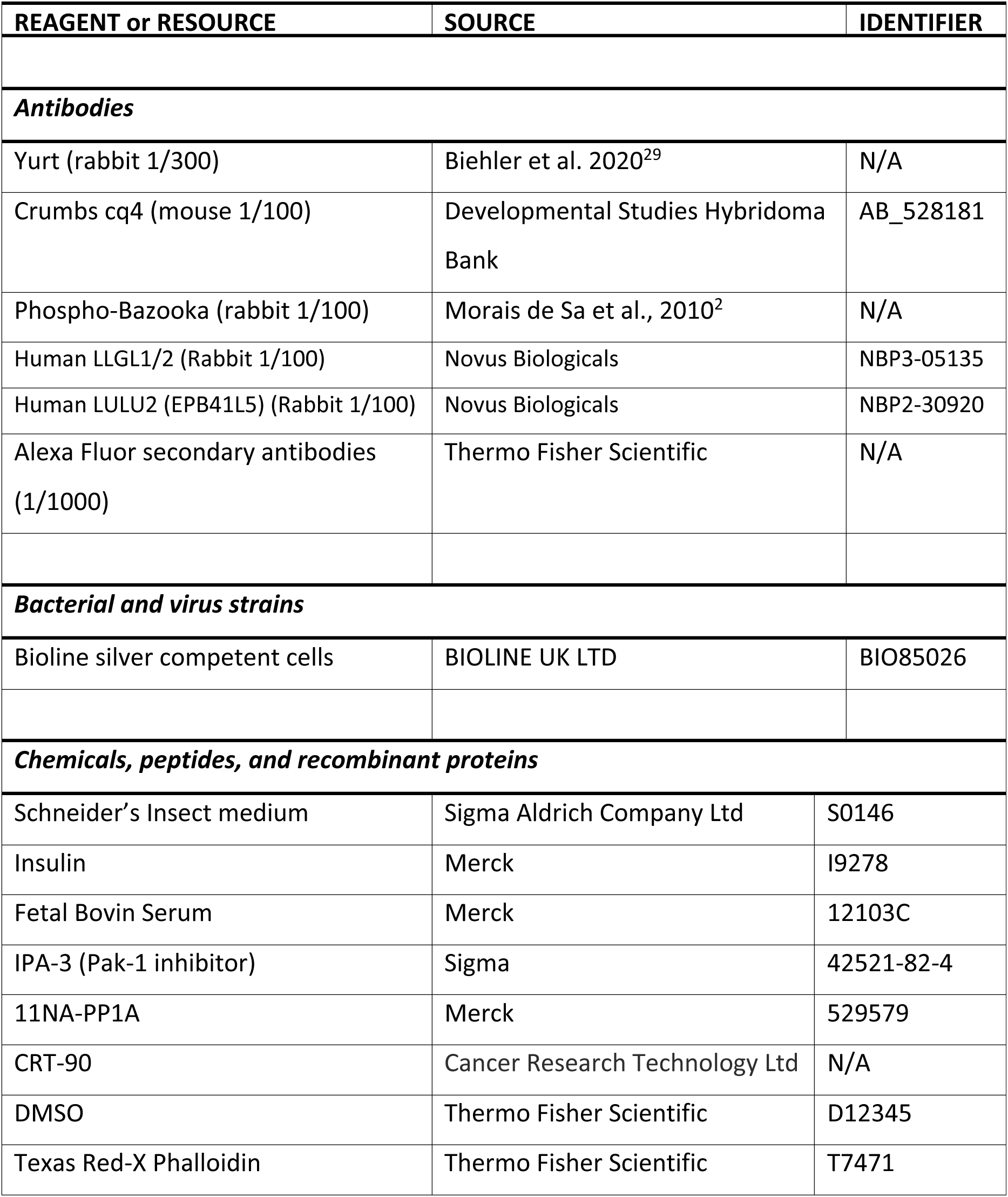
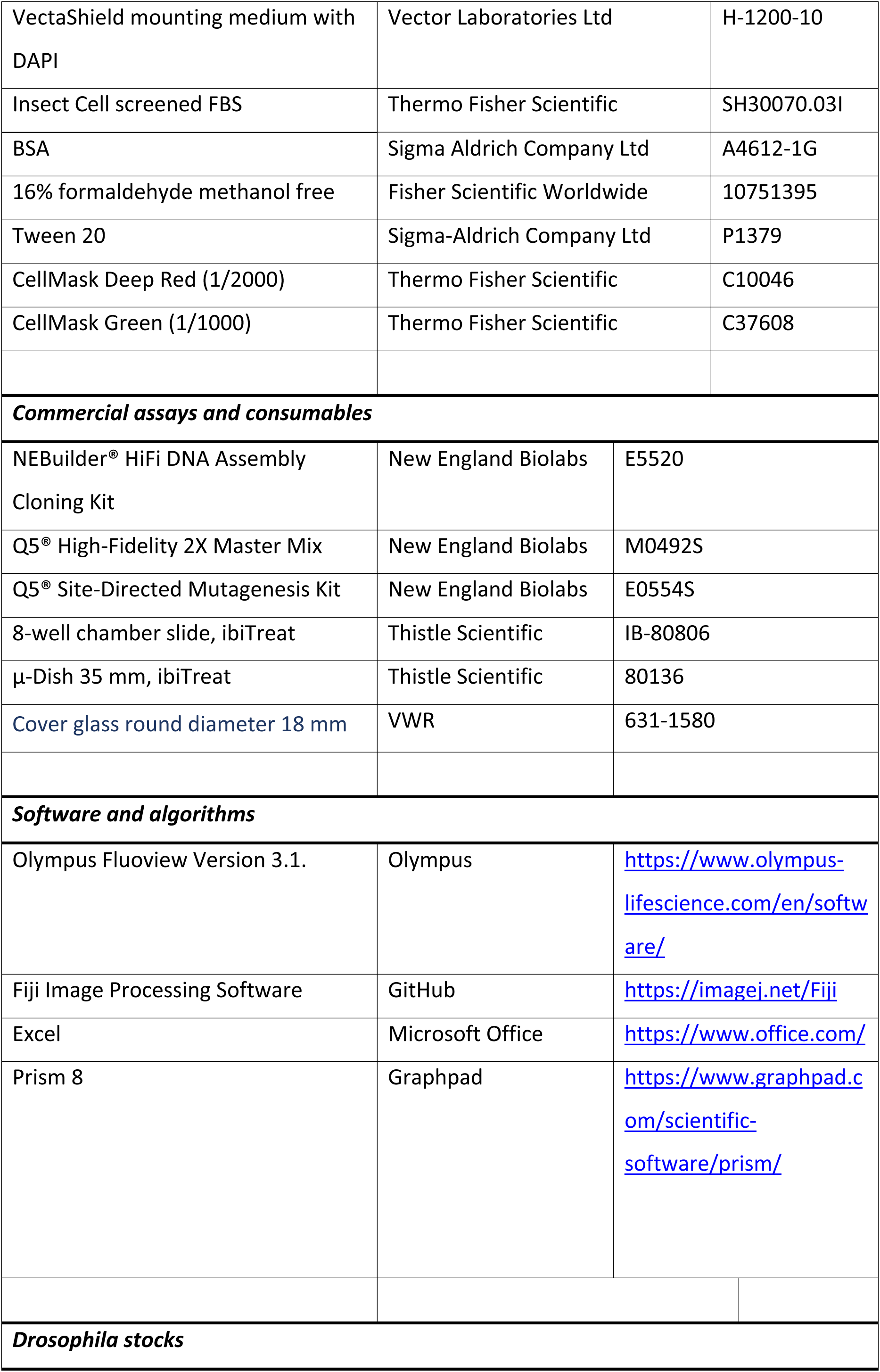
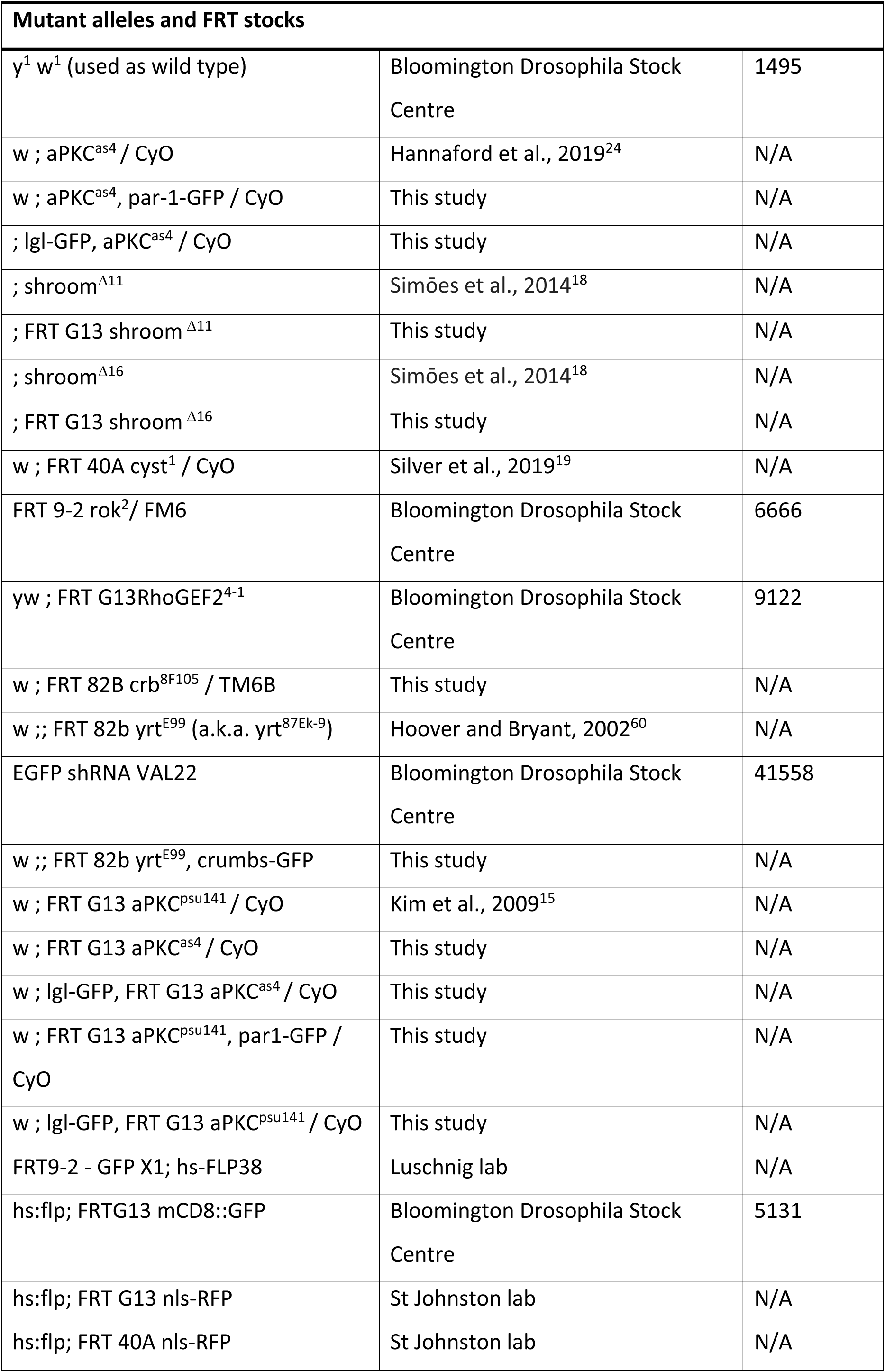
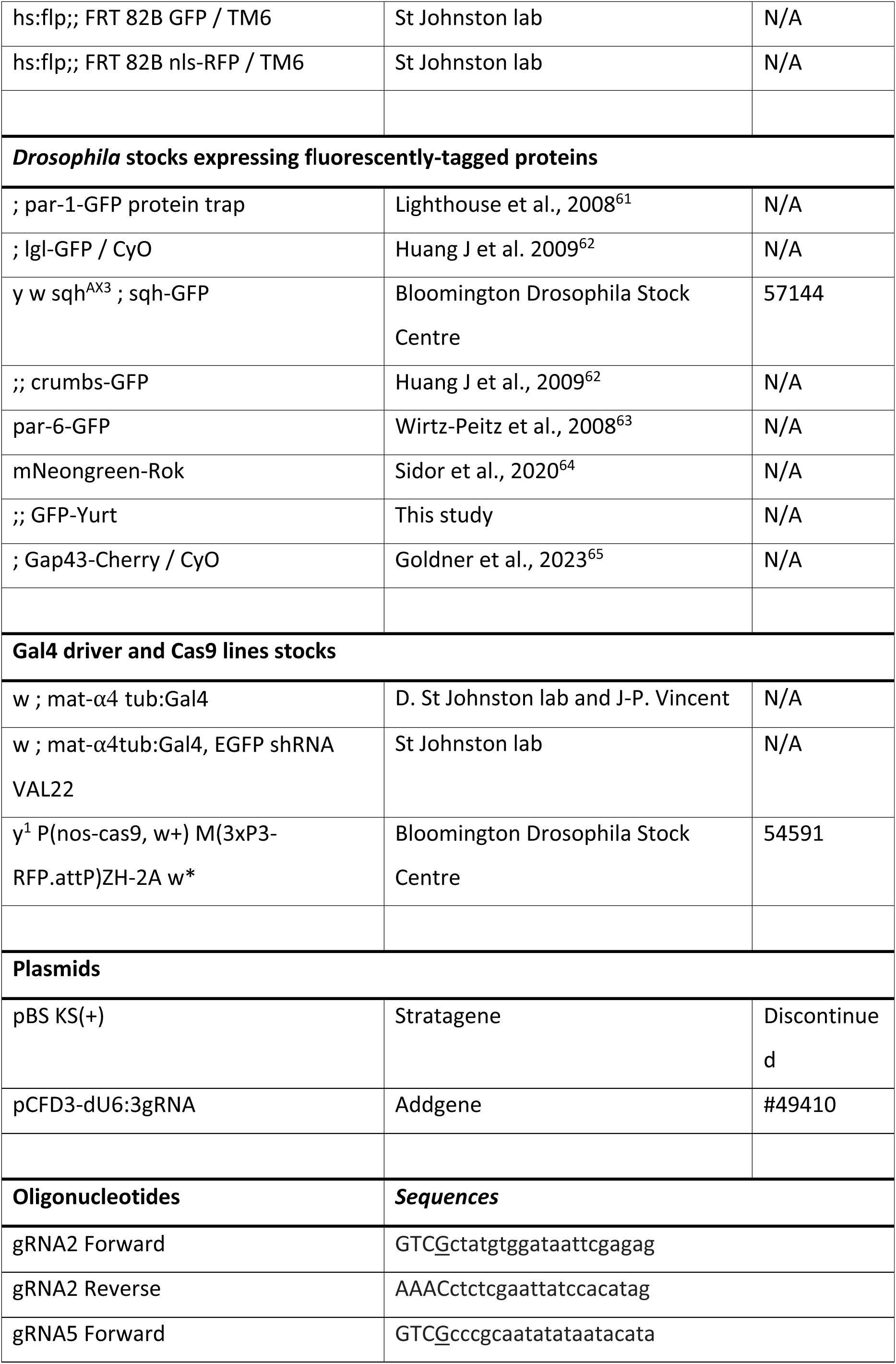
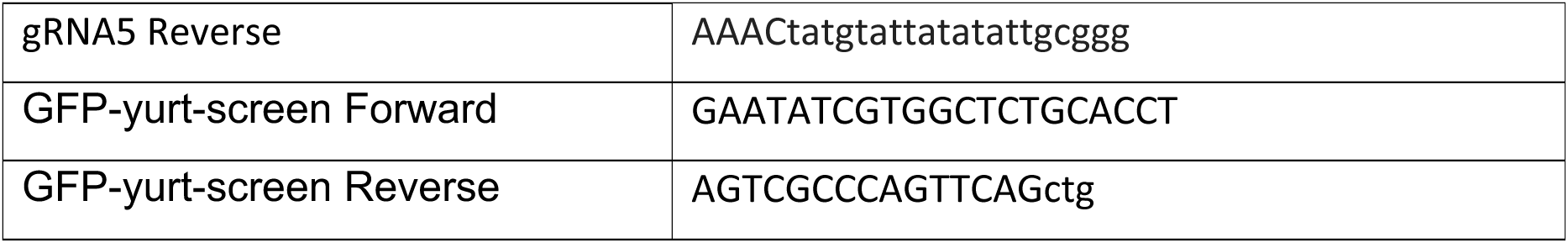

### Methods

#### Stock maintenance and *Drosophila* genetics

Standard procedures were used for *Drosophila* husbandry and experiments. Flies were reared on standard fly food supplemented with dried yeast at 25 °C. Heat shocks to induce germline clones were performed on Larvae stage 3 at 37 °C for 1 h (twice daily) for three days. Flies were kept at 25 °C for at least 3 to 5 days after the last heat shock before dissection. UAS-transgenes were expressed using Gal4 drivers in flies raised at 25°C; adult females were dissected 3 to 5 days after hatching.

#### Ovary dissection and drug treatment

Ovaries were dissected in Schneider’s insect medium supplemented with insulin (1:2000) and 10% fetal bovine serum (FBS). When needed, CellMask (1/2000) was added to the dissection media. For fixed ovary observation, ovaries were incubated on a nutating motion shaker with the drug diluted in the dissection media according to the conditions and times mentioned in the figure legends. Ovaries were fixed for 10 min in 10% PFA and 0.2% Tween 20 in PBS. For live ovary observation, ovaries were transferred to a chamber slide well containing 200 μl of dissection media and allowed to adhere to the well for 10 min. The drug was then added by adding 100 μl of dissection media containing the drug before imaging started. IPA-3, CRT-90 and 11NA-PP1Astock solutions were diluted in DMSO to a 10 μM concentration, and aliquots were kept at -70°C. The control contains the same amount of DMSO as the corresponding experiment.

#### Culture of Caco-2 3D cysts

Caco-2 cells (Caco2, ATCC, HTB-37) were cultured in Petri dishes or flasks with DMEM/F12 (Gibco, Cat. No. 10565018) supplemented with 10% heat-inactivated FBS (Gibco, Cat. No. A5209502) and 1x Penicillin-Streptomycin (Gibco, Cat. No. 15140122) in an incubator set at 37°C with 5% CO2. The medium was changed every 2 to 3 days. Caco-2 cells were split every 15-20 days once they reached 80-90% confluency and maintained in 2D culture. For 3D culture, Caco-2 cells were first trypsinised and passed through a mesh to obtain single cells. The single cells were suspended in DMEM/F12 medium and seeded in 30 μL of 1:1 Matrigel and DMEM/F12 medium in a 12-well plate, with each well containing 4 Matrigel domes of about 1000 Caco-2 cells. Each well received 2 mL of DMEM/F12 medium, which was refreshed every 2-3 days.

#### Immunostaining

Ovaries were fixed for 15 min in 4% formaldehyde and 0.2% Tween 20 in PBS. For phospho-specific antibody immunostainings, a phosphatase inhibitor solution was added to the PBS 0,2% Tween 20 solution. 50X phosphatase inhibitor solution kept at -80°C: 0.105g NaF (Sigma S79209), 0.540g β-glycerophosphate (Sigma G9422), 0.092g Na_3_VO_4_ (Sigma 450243), 5.579g Sodium pyrophosphate decahydrate (Sigma S6422), qsp 50 ml dH2O.

Ovaries were then blocked in 10% BSA in PBS with 0.2% Tween 20 for at least 1 h at room temperature. Samples were incubated with primary antibodies overnight in PBS with 0.2% Tween 20 and 1% BSA at 4°C and were then washed three times in PBS-0.2% Tween 20 for 30 min. They were then incubated in secondary antibodies for at least 3 h in PBS-0.2% Tween 20 and washed 3 times before mounting in Vectashield containing DAPI.

Caco-2 3D cysts were washed twice with ice-cold PBS, treated with gentle cell dissociation reagent (Stemcell, 100-1077) on ice to release them by depolymerising the Matrigel, and fixed with 4% (vol/vol) formaldehyde for 30 minutes on ice. Post-fixation, cysts were washed in PBS twice, permeabilised with 0.5% (v/v) Triton X-100 (Sigma-Aldrich) in PBS for one hour at room temperature, and blocked with blocking buffer (5% BSA and 0.5% (v/v) Triton X-100 in PBS). The samples were incubated overnight at 4°C with primary antibodies, followed by secondary antibodies (Alexa Fluor 488 or 647-conjugated donkey anti-rabbit/mouse IgG) under light-protected conditions at 4°C overnight. For counterstaining, samples were incubated with 4’,6-diamidino-2-phenylindole dihydrochloride (DAPI) (1 μg/mL; Fisher Scientific). The concentrations of primary antibodies used are indicated in the Key Resources Table. Secondary antibodies and Phalloidin were used at 1/500.

#### Egg chamber compression

Ovaries were dissected in Schneider’s insect medium supplemented with insulin (1:2000), 10% foetal bovine serum (FBS), and CellMask. Egg chambers beyond stage 9 were removed. The remaining egg chambers were transferred to a 35 mm ibiTreat dish containing 40 μL of dissection medium and allowed to adhere for 10 minutes. To maintain humidity, a piece of moistened Whatman paper was placed along the edge of the dish. Egg chambers were then covered with an 18 mm round coverslip, creating a suction effect between the dish and coverslip and applying gentle pressure to the samples prior to imaging.

### Imaging

Imaging was performed using an Olympus IX81 (40×/1.3 UPlan FLN Oil or 60×/1.35 UPlanSApo Oil).

Samples were imaged using an inverted Olympus IX81 (40× 1.35 NA Oil UPlanSApo, 60× 1.35 NA Oil UPlanSApo, 100x 1.35 NA Oil UPlanSApo) using the Olympus FluoView software Version 3.1 and processed with Fiji.

### N-terminal tagging of endogenous Yurt with GFP

The GFP-tagged Yurt line was generated via homology-directed repair using the Cas9-CRISPR system. The donor plasmid for homology-directed repair was constructed by inserting three fragments into the pBS KS(+) plasmid using NEB HiFi assembly: a 710 bp left homology arm corresponding to the genomic sequence upstream of the *yurt* ATG, a 714 bp GFP sequence in-frame with the *yurt* ATG, and the right homology arm downstream of the ATG. To prevent re-cutting, the PAM sites corresponding to both gRNA target regions were modified in the donor plasmid by introducing silent mutations using the Q5® Site-Directed Mutagenesis Kit. Two gRNAs targeting sequences near the *yurt* ATG were cloned into the pCFD3-dU6:3gRNA plasmid. The donor plasmid and both gRNA plasmids were injected at 150 ng/µl into CFD2 nos:Cas9 embryos. The GFP insertion upstream of the *yurt* ATG was screened by PCR using Q5® High-Fidelity 2X Master Mix and oligos located in the homology arms. PCR amplification yielded a 2871 bp fragment when GFP was inserted and a 2157 bp fragment when absent. The resulting GFP-tagged *yurt* line was confirmed to be homozygous viable and fertile. Oligos sequences are detailed in the Key Resource Table.

### Image Analysis

To measure the extent of proteins localisation to the apical domain of the follicle cells following aPKC inhibition, we calculated the ratio of the apical intensity over the lateral intensity. Using Fiji, we manually measured the mean intensity of the two follicle cell domains. The cytoplasmic background signal was measured and then subtracted it from both measurements. Measurements at the lateral side were divided by two, as they correspond to the lateral domains of two neighbouring cells.

All calculated ratios below 0 (due to subtraction of the cytoplasmic background) were normalised to 0.

### Statistical Analysis

All the statistical analyses were performed using Graphpad Prism software.

For single comparisons, Mann-Whitney unpaired t-test were performed for data that did not exhibit a normal distribution (Fig. 1).

For all other comparisons, a one-way non-parametric Anova was performed and a Kruskal-Wallis test was used to calculate significance between different conditions listed in figure legends.

## Resource availability

Lead contact: Further information and requests for resources and reagents should be directed to and will be fulfilled by the lead contact, Daniel St Johnston

Materials availability: All fly stocks generated in this study are available on request.

## Acknowledgements

We would like to thank Jens Januschke for sharing the aPKC^as4^ fly stock, and Ulrich Tepass, Jennifer Zallen and Eugenia Piddini for other fly stocks. We are very grateful to John Overton for injecting embryos to generate the N-terminally tagged GFP-Yurt stock and Richard Butler at the Gurdon Institute Imaging Facility for help with image analysis and quantification. CRT90 was a generous gift from Cancer Research UK Therapeutic Discovery Laboratories.

## Author contributions

DStJ and HD conceived and oversaw the project and analysed the data. HD, and AP designed and performed most experiments. NJ analysed the effects of *yurt* mutant clones after aPKC inhibition. CY generated the endogenously-tagged GFP-Yurt fly line. DStJ wrote the manuscript, which was revised and edited by all authors.

## Supplementary Material

**Supplementary Figure 1.**
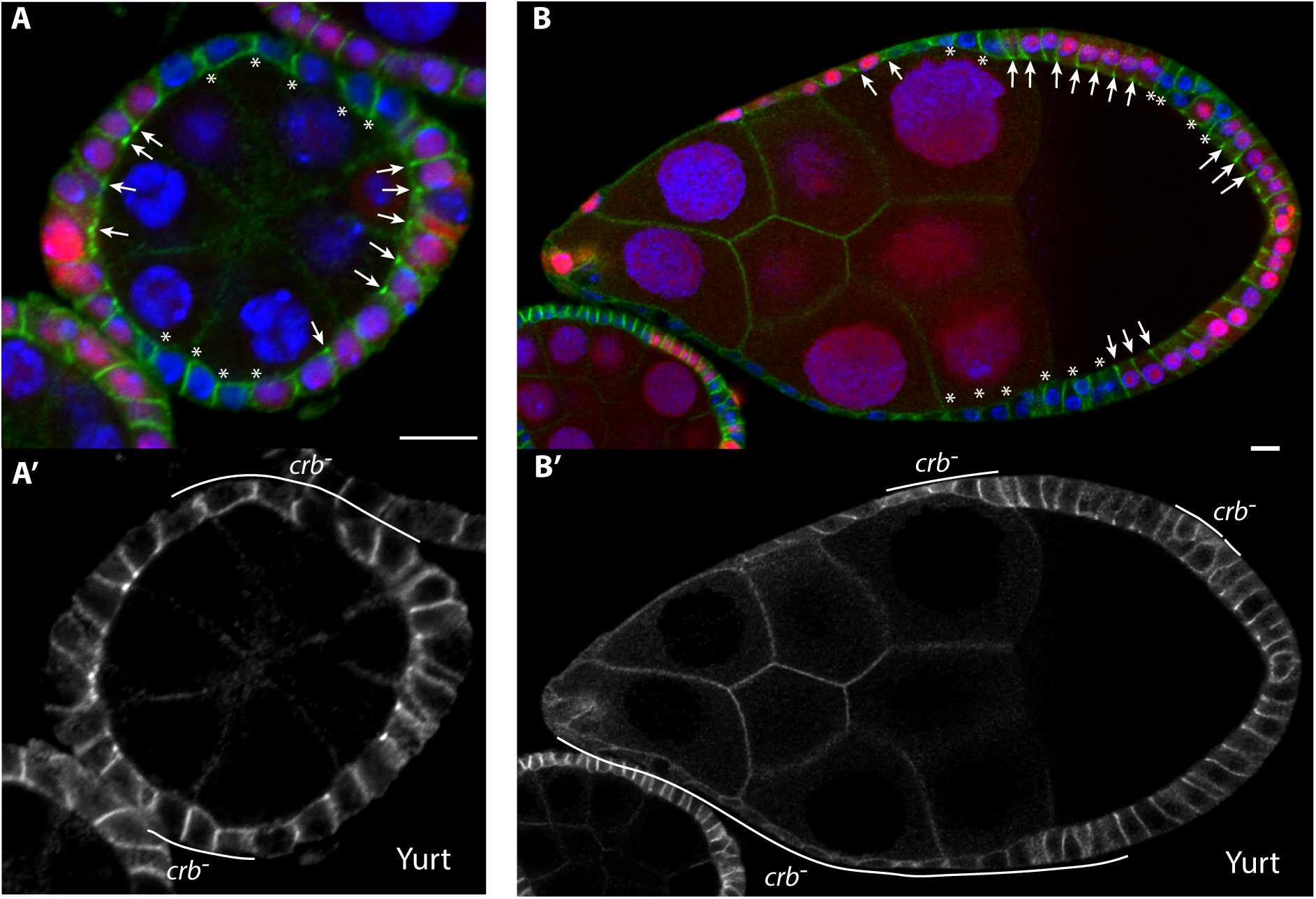
**A-B.** Untreated egg chambers expressing GFP-Yurt (green; greyscale in A’ and B’) containing *crb*^F105^ clones marked by the loss of nuclear RFP (red). Yurt is enriched at the apical junctions of the wildtype cells (arrowheads), but not at the apical junctions of *crb* mutant cells.

**Supplementary Video 1**. A GFP-Yurt expressing egg chamber treated with 10 µM CRT90. 1 frame/10 minutes for 30 minutes. Scale bar, 10 µm.

**Supplementary Video 2**. An *aPKC*^as^ homozygous egg chamber labelled with CellMask and treated with 1 µM 1NA-PP1A. 1 frame/minute for 10 minutes. Scale bar, 10 µm.

**Supplementary Video 3**. An egg chamber expressing MRLC-GFP to label Myosin, treated with 10 µM CRT90. 1 frame/minute for 12 minutes. Scale bar, 10 µm.

**Supplementary Video 4**. An egg chamber expressing Neon-Rok (green) containing *aPKC*^as^ clones marked by the loss of nls-RFP (red) and treated with 1 µM 1NA-PP1A. 1 frame/minute for 15 minutes. Scale bar, 10 µm.

**Supplementary video 5**. A Neon-Rok (green) expressing egg chamber containing *yurt*^E55^ clones marked by the loss of nls-RFP (red) treated with 10 µM CRT90. 1 frame/5 minutes for 40 minutes. Scale bar, 10 µm. The *yurt* mutant cells do not localise Rok apically or apically constrict and are instead stretched by the contraction of their neighbours.

**Supplementary video 6**. A GFP-Yurt (green) expressing egg chamber treated with 2 µM CRT90. 1 frame/ 5 minutes. Scale bar, 10 µm. Yurt localises to the apical Adherens junctions before spreading across the apical domain.

**Supplementary video 7**. A GFP-Yurt (greyscale) expressing egg chamber during compression. Yurt accumulates at the Adherens junctions and then spreads across the apical surface. 1 frame/ 5 minutes. Scale bar, 10 µm.

**Supplementary video 8**. Another GFP-Yurt (greyscale) expressing egg chamber during compression. 1 frame/ 5 minutes. Scale bar, 10 µm.

**Supplementary video 9**. Close up of a GFP-Yurt (greyscale) expressing egg chamber during compression, showing that yurt first localises to apical junctions before spreading across the entire apical domain. 1 frame/ 5 minutes. Scale bar, 10 µm.

